# Methods for simulations with thousands of interacting objects to model mass transport in the brain

**DOI:** 10.1101/2025.04.03.647052

**Authors:** Donald L. Elbert, Dylan P. Esguerra

**Author notes:** Corresponding author: Donald L. Elbert.

## Abstract

Clearance of toxic species that may cause neurodegenerative diseases relies on convection and diffusion of mass around cells in the central nervous system. In this manuscript, methods that allow the nanoscale modeling of mass transport around cells in the brain at a resolution of 8 nm are described. The cells are modeled as surface meshes and a parallelization scheme is used to solve the diffusion equation directly on the surface mesh of each biological cell independently. This is followed by mass exchange between biological cells that are in direct contact. The analysis volume size is fixed at 4 μm x 4 μm x 4 μm but arrays of analysis volumes of arbitrary size may be analyzed, with fluxes across analysis volume boundaries updated at each time step by a semi-implicit formulation. Setup of the discretized equations is described, along with the ‘face matching’ that allows mass transfer between cells. Parallelization is via a manager/worker framework. One-sided RMA communication in MPI is used to coordinate the efforts of multiple workers, with common information handled by the manager. This specialized framework is suitable for analyzing diffusional transport around thousands of interacting objects simultaneously. The framework is implemented with custom code developed in Julia, which is used here as a rapid prototyping language. Using model geometries, the accuracy of the discretization methods are demonstrated, with limitations and next steps described.

**Author’s summary:** The methods that enable modeling diffusional mass transport around thousands of objects in parallel are described. The accuracy of the method with model objects is demonstrated.

## Introduction

Solution of partial differential equations on 2D or 3D meshes may be performed with any number of available software tools. Commonly used platforms include the Ansys suite and OpenFoam. Additionally, there are over 11,000 ‘CFD’ repositories available on GitHub, with 101 in the Julia language alone. One reason for so many options is that the solution of the discretized equations is usually straightforward. However, the setup of the equations is complicated by processing complex mesh architectures and application of boundary conditions.

Most of the new framework is inspired by OpenFoam,^1^ but specialized for this problem. Typical CFD packages model flow around or inside one object with complex boundary conditions (obviously exceptions exist). Additionally, reaction-diffusion within realistic 3D models of neurons can be performed in the package Neuron,^2^ but the focus in this work is on the extracellular space. Here, there are thousands of objects and relatively simple and predictable boundary conditions, but with the mass transport occurring in the narrow gaps between the objects. While it is theoretically possible to adapt existing CFD packages to the current problem, adaptation could be more time consuming than rapid prototyping a purpose-built CFD code. Except for the Marching Cubes algorithm, all analysis code is written in Julia.^3^

The source of the imaging data used to construct the surface meshes is the Allen Institute’s Microns Explorer repository.^4–7^ The current project is focused on a small region of visual cortex, to demonstrate the capabilities of the approach. Because the TEM images resolve almost none of the extracellular space, a virtual space between adjacent cells is defined. This could be handled in several different ways, but the current approach is to assume a very thin and negligible layer of extracellular fluid surrounding surface meshes. Such an approach has been used to calculate geodesic distances by solving the heat equation directly on surface meshes.^8,9^ This is an approximate method but can yield insights about regions within the cortical tissue that may have higher amounts of toxic species, which may contribute to the development of neurodegenerative diseases.^10^ The current methods-focused paper is a companion to another paper describing biologically focused results.

## Methods

### Program: ImageSegmentation_14.0.py

1) Segmented images were downloaded from the Microns Explorer website (https://www.microns-explorer.org/cortical-mm3), centered at the location X = 340,796, Y = 122,482 (in 4 nm coordinates) or X = 170398, Y = 61241 (in 8 nm coordinates) and spanning Z = 17391-17591. Downloaded segmented image “frames” were 501 x 501 voxels (8 nm per voxel). The frames were stored with the following file name convention: frame_{zloc}_{xloc}_{yloc}_{height}.csv where zloc is the starting frame of the block, xloc and yloc are the center voxels of the block in 8 nm coordinates. Height is the length of a side of the analysis volume in microns, which in this case is 4 μm, corresponding to 100 frames per block due to the nominal slice thickness of 40 nm. The analysis volumes used in a simulation are specified in the file ‘Simulation_settings _4.0_smoothed.csv’ with the following table columns: Simulation number, xloc (x location of the center of the base analysis volume), yloc (y location of the center of the base analysis volume), zloc (starting frame z location), height, minus_x (number of analysis volumes in the -x direction), plus_x (number of analysis volumes in the +x direction), minus_y (number of analysis volumes in the -y direction), plus_y (number of analysis volumes in the +y direction), minus_z (number of analysis volumes in the -z direction), plus_z (number of analysis volumes in the +z direction), filled (0 if z frames have not yet been interpolated, 1 following interpolation)

### Programs: See companion publication for image processing programs

2) Likely segmentation errors were removed by mode filtering as described in a second companion publication. The smoothed frames were stored with the following file name convention: sframe_{zloc}_{xloc}_{yloc}_{height}.csv otherwise following the naming convention in step 1.

3) The resolution in X and Y directions was 8 nm, while the spacing between layers in the Z direction was nominally 40 nm. Rather than downsampling in the X and Y directions (which may cause fragmentation of dendritic spines), a new set of algorithms was used to interpolate between frames in the Z direction. After introducing four interpolated frames per original frame, a new zloc convention was adopted, which is simply 5x the original zloc. The zframes were mode filtered again, but this time were filtered in the xy plane, then the zx plane, then the yz plane, then the xy plane again. The smoothed zframes are stored as szframe_{zloc}_{xloc}_{yloc}_{height}.csv, with the new zloc (5x) convention.

4) The frame size of 501 x 501 voxels gives a one row or column of overlap between adjacent analysis volumes. However, the smoothing protocols currently yield different results on either side of the overlap region. When applicable, adjacent analysis volumes are stitched together by overwriting the boundary face of a volume with the boundary face of the adjacent volume in the positive x, y or z direction. The modified frames are stored as: szmframe_{zloc}_{xloc}_{yloc}_{height}.csv, with the zloc (5x) convention

### Program: unique_neurons_4.1_smooth.jl

5) The unique cells in an analysis volume are identified by the program unique_neurons_4.1_smooth.jl. The list of unique cells is sorted by size in reverse order and stored in frame_uniq_{zloc}_{end_zloc}_}_{xloc}_{yloc}_{height}.csv. ‘end_zloc’ is the last zloc in the stack. A file is also generated listing the coordinates of all the analysis volumes, ‘MarchingCubes_settings_smooth.csv’ from the data in ‘Simulation_settings _4.0_smoothed.csv’.

### Program: MarchingCubes3D_2.0a_smoothz_mpi.c

6) The Lewiner Marching cubes algorithm is applied to the smoothed and interpolated frames for each unique cell, exporting surface meshes as: {cell_ID}_{zloc}_{xloc}_{yloc}_{height}.ply.^11^ The ‘cell_ID’ is the 18-digit identifier from the MicronsExplorer dataset. A manager/worker MPI model was added to the existing code to produce multiple meshes in parallel.^12^

### Program: PlyToJLD2_3D_3.0a.jl

7) The ply files are analyzed in a Julia program to record nodes (vertices), faces, edges and boundary edges. The locations of the face centers are also recorded. The connectivity of the mesh is improved using the reverse Cuthill-McGee algorithm (SymRCM.jl). Meshes are stored in the following structure:

struct Structure_cell
npf::Int32 # Number of nodes per face
num_faces::Int32 # Total number of faces
num_nodes::Int32 # Total number of nodes
num_edges::Int32 # Total number of edges
num_boundary_edges::Int32 # Total number of boundary edges
faces::Array{Int32,2} # 12 x num_faces array
nodes::Array{Float32,2} # 3 x num_nodes array
normals::Array{Float32,2} # Surface normal for each face, 3 x num_faces array
edges::Array{Int32,2} # 11 x num_edges array
face_center::Array{Float32,2} # Face center location for each face, 3 x num_faces array
end

The arrays of nodes, faces and edges allow fast lookup and iteration over array elements. A structure is generated for each cell and saved as: {cell_ID}_{zloc}_{xloc}_{yloc}_{height}_structure.jld2. The JLD2 format is based on the HDF5 format and provides efficient compression automatically.

### File: simulation_settings.toml

8) Simulation settings are stored in a TOML file, including boundary conditions, source terms, initial conditions, time step size, diffusion coefficient, density and flow velocity. The boundary conditions are only applied at the exterior boundary of an array of analysis volumes. In future versions of the program, boundary conditions will also be applied at the surfaces of capillaries. The TOML file settings are read into the structure Simulation Parameters.

struct Simulation_parameters
Δt::Float32
BCarray::Vector{BC}
ICvalue::Float32
ℾ::Float32
ρ::Float32
ū::Vector{Float32}
Source_u_::Float32
Source_p_::Float32
End

The boundary conditions are stored in BCarray as a vector of BC structures:

struct BC
ID::Int32
type::Int32
value::Float32
end

where ID is the number of the boundary or patch, type is Dirichlet (‘1’), Neumann (‘2’) or interior (‘3’) (i.e. interior to an array of analysis volumes).

### Program: neuron_transport_setup_3D_3.0a.jl

9) The surface mesh described in ‘Structure_cell’ is further analyzed to calculate: a) weights of adjacent nodes for calculation of nodal concentrations, b) non-orthogonality correction using the over-relaxed approach, c) face ‘volume’, d) edge normal vector, e) edge ‘area’, f) center-to-center vector from over-relaxed approach, g) center-to-center magnitude from over-relaxed approach, h) edge tangent vector by over-relaxed approach i) distance between adjacent face centers, j) face center-to-face center vector, k) geometric weighting factor for each edge (inverse of distance from face center to edge midpoint).

struct Solution_setup
Sim_parameters::Simulation_parameters
weight_node::Array{Float32,2} # weights of adjacent nodes for calculation of nodal concentrations t^_dot_lf_δ_edge::Array{Float32,2} # non-orthogonality correction using the over-relaxed approach volume_face::Vector{Float32} # face ‘volume’
S::Array{Float32,2} # edge normal vector
Sedge::Array{Float32,3} # edge ‘area’
E::Array{Float32,2} # center-to-center vector from over-relaxed approach
Ef::Vector{Float32} # magnitude of E
T::Array{Float32,2} # edge tangent vector by over-relaxed approach
dCF::Vector{Float32} # distance between adjacent face centers
rCF::Array{Float32,2} # vector between adjacent face centers
weight_edge::Vector{Float32} # geometric weighting factor for each edge (inverse of distance from face center to edge)
end

A structure is generated for each cell and saved as: {cell_ID}_{zloc}_{xloc}_{yloc}_{height}_setup.jld2.

10) Each biological cell is then compared with all other biological cells in an analysis volume for face overlap. Face overlap is detected by overlap of face centroids. The matches are recorded by pairing the matched face numbers for the two cells. To prevent double storage, the biological cell that is earlier in the list of cells within an analysis volume is designated as the owner of the match, and the paired cell is designated the neighbor. The structure ‘Match_face’ contains an array ‘match_faces’ with the ‘face numbers’ of matches, the total number of face matches (needed for pre-allocation), a local ‘owner’ array for each cell (‘owner_temp’), and the total number of cells that match with other cells (needed for pre-allocation).

struct Match_face
match_faces::Matrix{Int32}
total_face_matches::Int64
wner_temp::Matrix{Int32}
number_matched_cells::Int32
end

A structure is generated for each cell and saved as: {cell_ID}_{zloc}_{xloc}_{yloc}_{height}_match.jld2.

11) The ‘owner_temp’ arrays for all the cells in the analysis volume are then consolidated into a single ‘owner’ array for the entire analysis volume. The ‘owner’ array is then sorted by the neighbor cell numbers to produce the ‘neighbor’ array. A map of pointers into ‘owner’ and ‘neighbor’ based on cell numbers is recorded in ‘owner_neighbor_ptr’. These arrays are stored in the structure ‘Owner_neighbor’, along with information that allows pre-allocation later, such as the number of cells with matches (‘num_cells’), the total number of matched faces (‘num_matches’), a vector of the total number of faces matched for each cell (‘num_faces_array’), and the list of 18-digit cell IDs present in the volume (‘cell_list’).

struct Owner_neighbor
num_cells::Int32
num_matches::Int64
cell_list::Vector{Int64}
wner::Matrix{Int32}
neighbor::Matrix{Int32}
neighbor_sort_vec::Vector{Int32}
wner_neighbor_ptr::Matrix{Int32}
num_faces_array::Vector{Int32}
end
A structure is generated for each analysis volume and saved as: simulation_{zloc}_{xloc}_{yloc}_{height}.jld2.

12) Each boundary edge is identified by both nodes of the edge being located on the boundary (within a certain precision). Edges that are on the boundary of an analysis volume are written into the array ‘boundary edges’, which is sorted by boundary type (exterior versus interior faces of an analysis volume). If the boundary edge is on an interior face in an array of analysis volumes, the relevant elements of the struct ‘Structure_cell’ are recalculated with respect to the adjacent face in the neighboring analysis volume. The list of faces on interior boundaries is stored in an ‘interior_boundary_faces’ vector for fast lookup of ‘ghost’ concentration values at the end of each time step. The number of boundary edges, a pointer to the start of interior boundary edges in ‘boundary edges’, a map of edge numbers to boundary edge numbers and a list of interior boundary nodes are stored along with ‘boundary_edges’ and ‘interior_boundary_faces’ in the structure ‘Boundary_edges’.

struct Boundary_edges
boundary_edges::Array{Int32,2} # number of boundary edges
interior_boundary_faces::Array{Int32,1}
sum_interior_faces::Int32 # total number of faces at interior boundaries
split_boundary_edges::Int64 # pointer to the start of interior boundary edges in ‘boundary edges’
edge_to_boundary_edge::Vector{Int64} # map of edge numbers to boundary edge numbers
interior_boundary_nodes::Array{Int32,2}
end

A structure is generated for each cell and saved as:

{cell_ID}_{zloc}_{xloc}_{yloc}_{height}_boundary_edges.jld2.

13) The setup program then records additional information about interior boundary edges. The number of edges of each cell that lie on each of the six faces of the analysis volume is recorded (‘cells_on_faces’). This array also stores pointers to the start of that boundary’s starting position in ‘interior_boundary_faces’. Finally, this array also stores pointers to the start of that cell’s starting position in the vectors ‘cube_face_conc’ and ‘cube_face_conc_ghost’. These vectors store concentration values for interior boundary faces for the entire analysis volume face. Finally, for pre-allocation, the total number of cells that pass through each face of an analysis volume is recorded in ‘cells_on_cube_side’, while the total number of edges passing through the analysis volumes faces is recorded in ‘edges_on_cube_side’. The former is simply the sum of Booleans for the test ‘>0’ on each element in the first six rows. The latter is simply the sum of each of the first six rows of ‘cells_on_faces’. These arrays are stored in the structures ‘Cube_properties’ and ‘Ghost_cells_dynamic’.

struct Cube_properties
cells_on_faces::Array{Int32,2}
cells_on_cube_side::Vector{Int32} edges_on_cube_side::Vector{Int32}
end

A structure is generated for each analysis volume and saved as: frame_uniq_{starting_zloc}_{ending_zloc}_{xloc}_{yloc}_{height}.jld2.

where starting_zloc is the first z-frame and the ending_zloc is the last z-frame.

struct Ghost_cells_dynamic
cube_face_conc::Vector{Vector{Float32}}
cube_face_conc_ghost::Vector{Vector{Float32}}
end

A structure is generated for each analysis volume and saved as: frame_uniq_{starting_zloc} _{ending_zloc}_{xloc}_{yloc}_{height}_ghost_dynamic.jld2.

14) Calculation of the implicit and explicit parts of the flux across analysis volumes requires sharing of structural information of ‘ghost’ cells amongst neighboring analysis volumes. Unlike ‘Ghost_cells_dynamic’, the structure ‘Ghost_cells_static’ contains information that does not need to be updated. ‘Ghost_cells_static’ is similar to ‘Structure_cell’ but records information only for internal boundary faces and their ghost face counterparts.

struct Ghost_cells_static
cube_face_S::Vector{Array{Float32,2}}
cube_face_E::Vector{Array{Float32,2}}
cube_face_Ef::Vector{Vector{Float32}}
cube_face_T::Vector{Array{Float32,2}}
cube_face_dCF::Vector{Vector{Float32}}
cube_face_rCF::Vector{Array{Float32,2}}
cube_face_weight_edge::Vector{Vector{Float32}}
cube_face_t^_dot_lf_δ_edge::Vector{Vector{Float32}}
end

A structure is generated for each analysis volume and saved as: frame_uniq_{starting_zloc} _{ending_zloc}_{xloc}_{yloc}_{height}_ghost_dynamic.jld2.

15) A sparse representation of the coefficient matrix *a_i,j_* (i.e. the **A** matrix in the linearization **Aϕ** = **b**) is pre-allocated and stored, where **ϕ** is the concentration at each face. In the current framework, there is no need to update this matrix during analysis. The boundary condition and source term contributions to the non-homogenous terms (**b**) are recorded in the vector ‘source_u_’. The nodal values are fixed to the boundary conditions at exterior faces of analysis in the array ‘boundary_node_values’. All of these are collected and put into the structure ‘Matrix_setup’.

struct Matrix_setup
source_u_::Vector{Float32}
a_sparse::SparseMatrixCSC{Float32, Int32}
boundary_node_values::Array{Float32,2}
end

A structure is generated for each cell and saved as:

{cell_ID}_{zloc}_{xloc}_{yloc}_{height}_matrix_setup.jld2.

### Program: neuron_transport_solve_3D_3.0a.jl

16) At the first time step, the initial values are applied to all faces, including the ‘ghost cells’ of interior boundary edges.

17) At each time step for each biological cell, the sparse **A** matrix is read from disk, the **b** vector is constructed from source_u_, the unsteady term, and the non-orthogonality correction. The non-orthogonality correction is calculated by estimating the tangential gradient from the nodal values, and the correction is updated at each iteration of the solver. The solution of the linear system is by the conjugate gradient method and stored in vector ‘phi’. Concentration values at matched faces are recorded in vectors ‘phi_owner’ or ‘phi_neighbor’, as appropriate and as illustrated in the results section. Concentration values at interior boundary faces are recorded in the vector ‘cube_face_conc’.

18) At the end of each time step, the concentrations at matched faces are averaged between adjacent cells and ‘phi’ for each cell is updated with the averaged values. The concentrations at interior boundary edge faces in ‘cube_face_conc’ are written into ‘cube_face_conc_ghost’ in the neighbor analysis volume. For visualization, results are stored in the vtk format.

## Results

### Overall approach

The greatest challenges are: 1) A reasonable size for ‘analysis volumes’. The analysis volume size of 4 μm x 4 μm x 4 μm is largely dictated by the memory requirements of the current Marching Cubes algorithm. The ultimate goal is to analyze an entire capillary bed consisting of 32,768 analysis volumes. Parallelization by analysis volumes is possible but the current approach focuses on parallelizing by biological cell within analysis volumes and the number of available cores is typically less than the number of biological cells. 2) The cost/benefit of parallelizing by biological cell versus fusing all meshes in an analysis volume is still to be determined. 3) This proof-of-concept project models diffusion directly on surface meshes. While this is common practice to determine geodesics by the ‘heat method’,^8,9^ the accuracy of the method decreases as the angles between faces are <<180°. Inflating the surface meshes to produce prism layers will improve accuracy, but the smoothness of the mesh needs to be high enough to prevent self-intersections and negative volumes. Progress in modeling of convection/diffusion/reaction and pressure-velocity coupling is tightly coupled to progress in producing smooth but accurate meshes. 4) The extracellular space is not well-preserved in the TEM images and the best way to model the architecture of the extracellular space is to be determined. 5) Co-registration of neuron somas in TEM images with optical two-photon images showed an 8% decrease in the x-z plane (parallel to the surface of the brain) and a 14% expansion in the y-direction (dorsal-ventral axis).^7^ This results in a negligible net volume change, yet the extracellular matrix is practically absent in the TEM images when compared to cryo-ET performed without a dehydration step.^13^ Even with these limitations, the current approach is able to model diffusion of a solute out of the brain cortex with 2175 biological cells and 313 million face elements, as described in a companion paper. The current paper is focused on the data structures used, the face matching procedure and the ghost cell formulation.

### Node, edge and face storage

As shown in Figure 1, nodes (vertices) of the triangular surface mesh are stored with x, y and z coordinates and each node is stored in one column, where the column number is implicitly the node number (Julia is column major, so reading columns is faster than reading rows).

**Figure 1:**
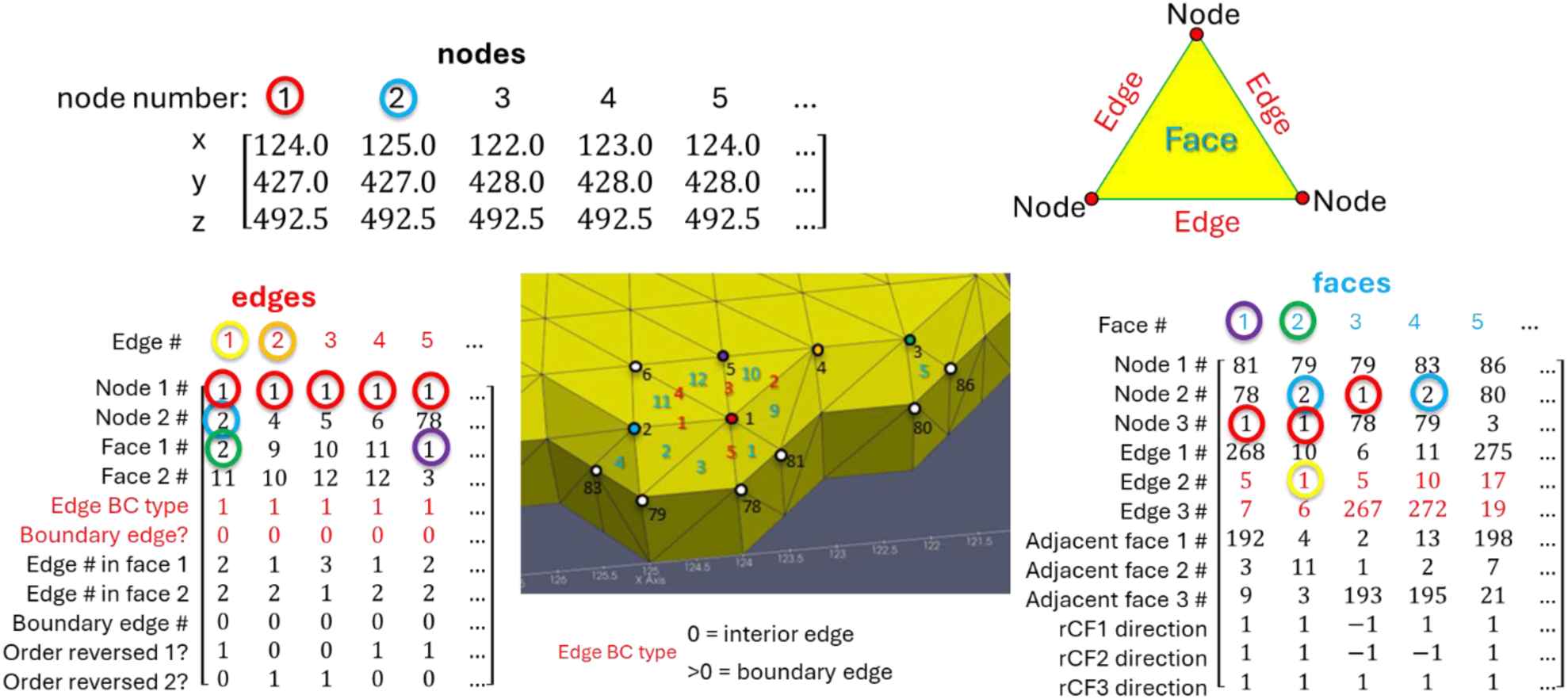
Data structures for nodes, edges and faces. “nodes” records the location of mesh vertices. “edges” stores the nodes connected by the edge and the faces adjacent to the edge. If the edge is on a boundary (on the face of an analysis volume), the BC type records if the edge is adjacent to another analysis volume and the edge’s number in the neighbor volume. The orientation of the edges with respect to the face normal is also recorded. “faces” records the adjacent nodes, edges and faces, and records the direction of the vector from the face center to adjacent face centers. Entries circled with the same color refer to the same node, edge or face.

Faces are stored in columns, with column number being an implicit representation of the face number. In most mesh formats, the number of vertices in a face is recorded followed by the node numbers of the vertices. In the present case, only triangular elements are used, obviating the need to store the number of vertices per face. Unlike traditional formats, the face array here carries along all the information that might be needed for present and many future solution methods. The size of the face array may be reduced in the future as the numerical method choices are solidified. The information stored in the face array for each face are: nodes, edges, adjacent faces and the direction of the vector with the adjacent faces. The face array is 12 x n_faces bytes, where n_faces is the number of faces in a cell’s surface mesh.

The edge array is 11 x n_edges bytes (n_edges is the number of edges). The edge array stores each edge’s nodes, faces, boundary condition type, boundary edge flag, edge number in first face and the second face, boundary edge number, and order of nodes in face 1 and in face 2. The edge array is only used in the setup phase, but ‘boundary_edges’, which is derived from the ‘edges’ array, is used at each iteration of the iterative solver. The ‘boundary_edges’ array records much of the same information as edges but also stores information about the analysis volume boundary upon which it resides, by changing the boundary condition type into a ‘boundary type’. The boundary type has a value of 1-6, these will refer to the face of the analysis volume upon which the edge lies, with 1 = -x face, 2 = +x face, 3 = -y face, etc. If the edge lies on an analysis volume face that is interior to the array of analysis volumes, the boundary type = 7. The boundary edge flag is also replaced with the analysis volume face number, which is -1 for exterior faces and otherwise follows the same convention as boundary type.

### Discretization of the convection/diffusion equation

Figure 2 shows an adaptation of a common 2D CFD approach and nomenclature. On 2D unstructured meshes the vector **S** is normal to each edge, with the magnitude of **S** being the ‘face’ area of the adjoining edge. In the current case, the length of the edge is a proxy for the face area of the prismatic volume that would result from inflation of the surface mesh. Here, **S** is a vector that is perpendicular to the edge *and* in the plane formed by ***d***_*CF*_ and *t̑*. Thus, ***S*** is calculated from the triple vector product ***n*** = *t̑* × (***d***_*CF*_ × *t̑*) with ***S*** = *area x **n***/‖***n***‖. The line connecting triangle centers (***d****_CF_*) is generally not parallel to ***S*** even in the 2D case (non-orthogonal) nor will it necessarily pass through the midpoint of the edge (skewness). A vector ***E*** in the direction of ***d****_CF_* is calculated as: ***E*** = (‖***S***‖/*cosθ*)***ȇ*** where ***ȇ*** is the unit vector in the direction of ***d****_CF_*. The concentration gradient the direction of **E** is **▽***ϕ* · ***ȇ*** = (*ϕ*_*F*_ − *ϕ*_*C*_)/‖***d***_*CF*_ ‖. A correction for the non-orthogonality correction between ***d****_CF_* and ***S*** is calculated using the over-relaxed approach.^1^ This correction requires the gradient in concentration at the edge midpoint. This is typically calculated using a Gauss-Green approach, which requires mesh inflation.^1^ Instead, the gradient is estimated by dividing the nodal concentrations by the length of the edge. This approach requires calculation of the nodal concentration, a calculation that most CFD codes attempt to avoid. However, it is possible to calculate the nodal concentrations rapidly by iterating through the columns of the data structures in Figure 1, using the recorded nodal weights and node numbers to sum the nodal concentrations. The entire calculation may be performed in one pass through the face array plus one pass through the array of boundary edges (the latter to fix external boundary conditions for nodes on the boundary). The boundary edges that are on exterior boundaries are first in the ‘boundary_edges’ array, such that only part of the ‘boundary_edges’ array needs to be iterated. The tangential vector **T** = **S** – **E** is parallel to the edge and its magnitude represents the strength of the cross-diffusion term. The non-orthogonal (cross-diffusion) correction is given by:

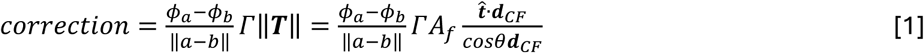

**Figure 2:**
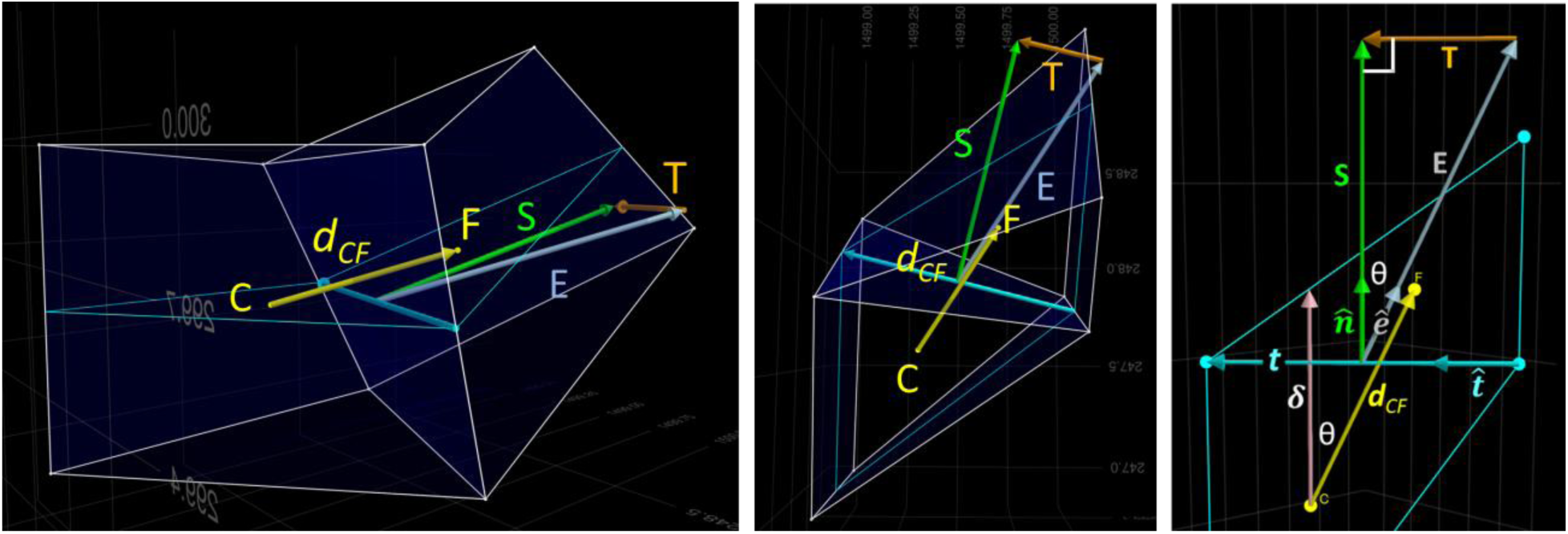
Views of two faces of a surface mesh connected by a common edge. Cyan triangles are the surface mesh. The white prisms are an inflation layer. ***d****_CF_* connects the centers of the cyan triangles (*C* & *F*). The ***S*** vector is perpendicular to the shared edge and in the plane formed by ***d***_*CF*_ and *t̑*, where *t̑* is the unit vector of the common edge. ***E*** is parallel to ***d****_CF_*. ***T*** is (***E*** – ***S***). The magnitude of ***S*** is the ‘area’ of the face, which here is the length of the shared edge. The magnitude of ***E*** is chosen such that ***T*** is parallel to the shared edge. **T** is used to correct for the angle between **S** and ***d****_CF_* (non-orthogonality correction). The third panel shows a case where the faces are coplanar. The angle between the face unit normal ***n̑*** and the unit normal in the direction of **E** (***ȇ***) is θ. The projection of ***d****_CF_* onto ***n̑*** is δ.

Where a and b are node locations associated with an edge, *ϕ* are concentrations, and Γ is the diffusion coefficient.^1,14^

The non-orthogonality correction is applied as a source term that adds to the non-homogenous **b** vector in **Ax** = **b**. The **A** matrix of coefficients of the discretization is filled and sparsified during problem setup and is not altered during the solution process. After addition of the explicit part of the unsteady term, the equations are solved by the conjugate gradient method, as the A matrix is symmetric and positive definite.

The offset between ***d_CF_*** and the edge midpoint is not corrected here (skewness correction). While convection terms are present in the discretization, they are only relevant when the mesh is aligned with the flow direction. Accounting for velocity on the curved domains would require solution of the Navier-Stokes equations, which requires accurate calculations of gradients and is dependent on ongoing work on surface inflation.

Results for solution of the convection/diffusion equation on model surface meshes is found in Figure 3. The solution on a rectangular 2D mesh was produced in Ansys Fluent (Figure 3A) and is compared to the solution on a 3D surface mesh of a square channel (Figure 3B). In Figure 3C, the solution is shown at the same time point but with diffusion only. The solution to the diffusion equation on the surface mesh of a circular pipe is shown in Figure 3D. The results from Figure 3B-D at different time points are shown in Figure 4 and compared with the solution of the 1D equation. In all cases, the accuracy of the method is high, demonstrating the correctness of the approach on these model geometries. Each of the model geometries consists of two objects joined in the middle. The motivation for fixing the right boundary condition at 1.0 is that a mass flux imbalance at the junction of the two objects would be masked if both boundary conditions were 0.0. With the current setup, a concentration gradient is present at the interface for most of the simulation.

**Figure 3:**
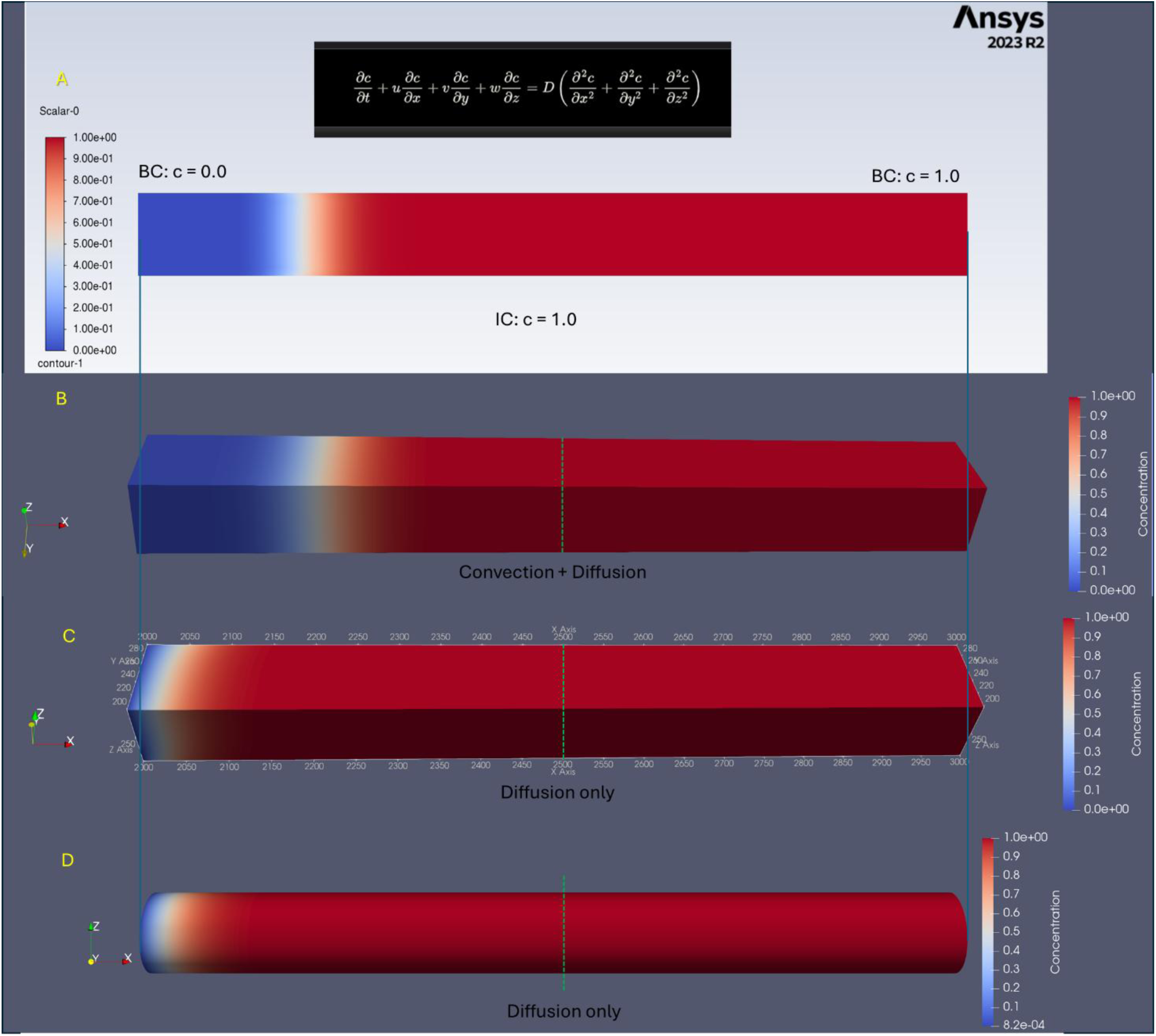
Validation of the surface mesh diffusion algorithm. Initial condition c = 1.0 at time = 0, boundary conditions concentration at left boundary = 0.0, concentration at right boundary = 1.0. Length = 1000 units (each unit = 8 nm). The diffusion coefficient of the solute is 6.4 x 10^-7^ cm^2^/s, typical for a peptide. Velocity = 0.16 cm/s, Peclet number = 200. Solutions shown at time point = 1 ms. A) Solution to the convection/diffusion equation (inset) on a 2D rectangular mesh in Fluent (Ansys 2023 R2), time step = 1 μs. B) Solution by the current method for convection and diffusion on the surface of a square channel, time step = 1 μs. C) Solution with velocity = 0.0 m/s (diffusion only). Time step = 10 μs. D) Solution with velocity = 0.0 cm/s on a circular pipe, time step = 10 μs. In panels B-D, the meshes and volumes are split at the midpoint and equations for each volume are solved sequentially, with mass flux calculated at the interface using a semi-implicit approach at the start of each time step.

**Figure 4:**
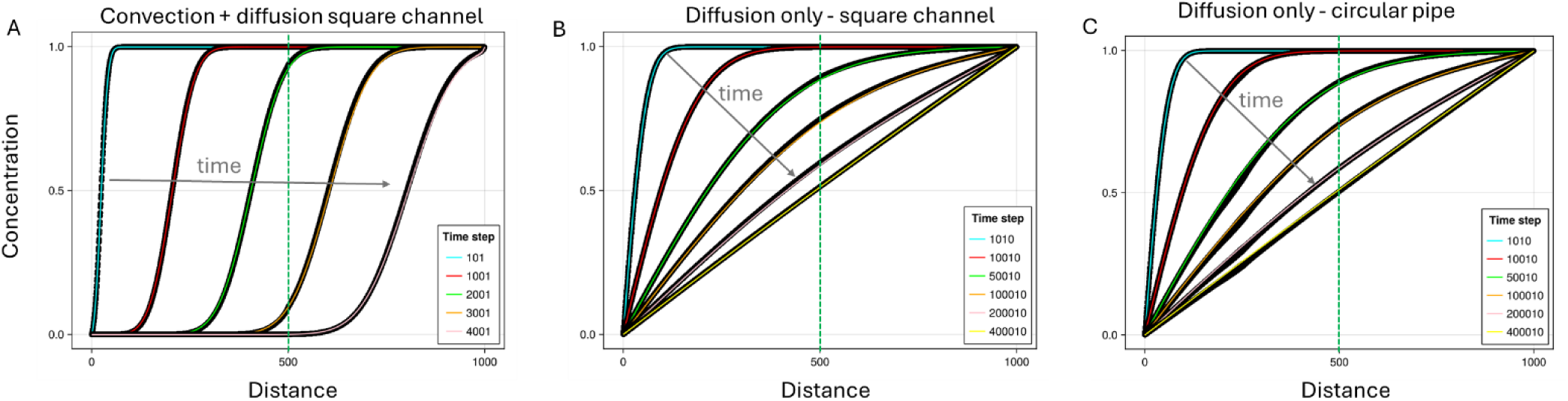
Agreement between 3D surface diffusion model and 1D solution. Solid lines are the method of lines solution (for convection/diffusion) or the exact solution (diffusion only). All faces in the square channel (total faces = 396,000) and circular pipe (total faces = 315,000) are plotted as black circles. A) Square channel convection + diffusion, B) Square channel, diffusion only, C) Circular pipe, diffusion only. Each distance unit = 8 nm. All other conditions are as described in the previous figure. The dashed green line is the interface between two analysis volumes.

### Face matching

To demonstrate the process of face matching, five cells are selected from the 431 cells in the base analysis volume (Figure 5A). The cell labeled “1” is in contact with cells 3 and 5, while cell 2 is in contact with cells 4 and 5, cell 3 is in contact with cells 1 and 5, cell 4 is in contact with cells 2 and 5, and cell 5 is in contact with all other cells (also see Figure 6). Contact is defined by the cell centroids of faces from different cells being identical to within a small tolerance. In these cases, not only do the cell centroids overlap, but all three of the vertices overlap, due to the details of the marching cubes algorithm. The shared faces between cell 5 and the other cells are shown in Figure 5B. Each cell in most cases shares faces with multiple cells, and all these matched faces are stored in only two vectors for each cell. One vector stores the face numbers for face matches that the cell ‘owns’ (‘owner_match’) and the other stores the face numbers for matches where it is the ‘neighbor’ (‘neighbor_match’), which are shown in Figure 7. A cell owns a match if its cell number is lower than its partner, where the cell number refers to the position of the cell in the list of the 18-digit identifiers (‘cell_list’). The array ‘owner’ (Figure 5C) is a map of all the matched pairs of cells and the location of the matched faces in ‘owner_match’. The array ‘neighbor’ stores essentially the same map but reordered by sorting row 2 and then by row 1 and additionally storing the location of the matched faces in the array ‘neighbor_match’ (Figure 5D). The array ‘owner_neighbor_ptr’ stores a map such that given a cell’s number, the relevant locations within owner and neighbor may be found along with the size of the vector needed to store the matches (Figure 5E).

**Figure 5:**
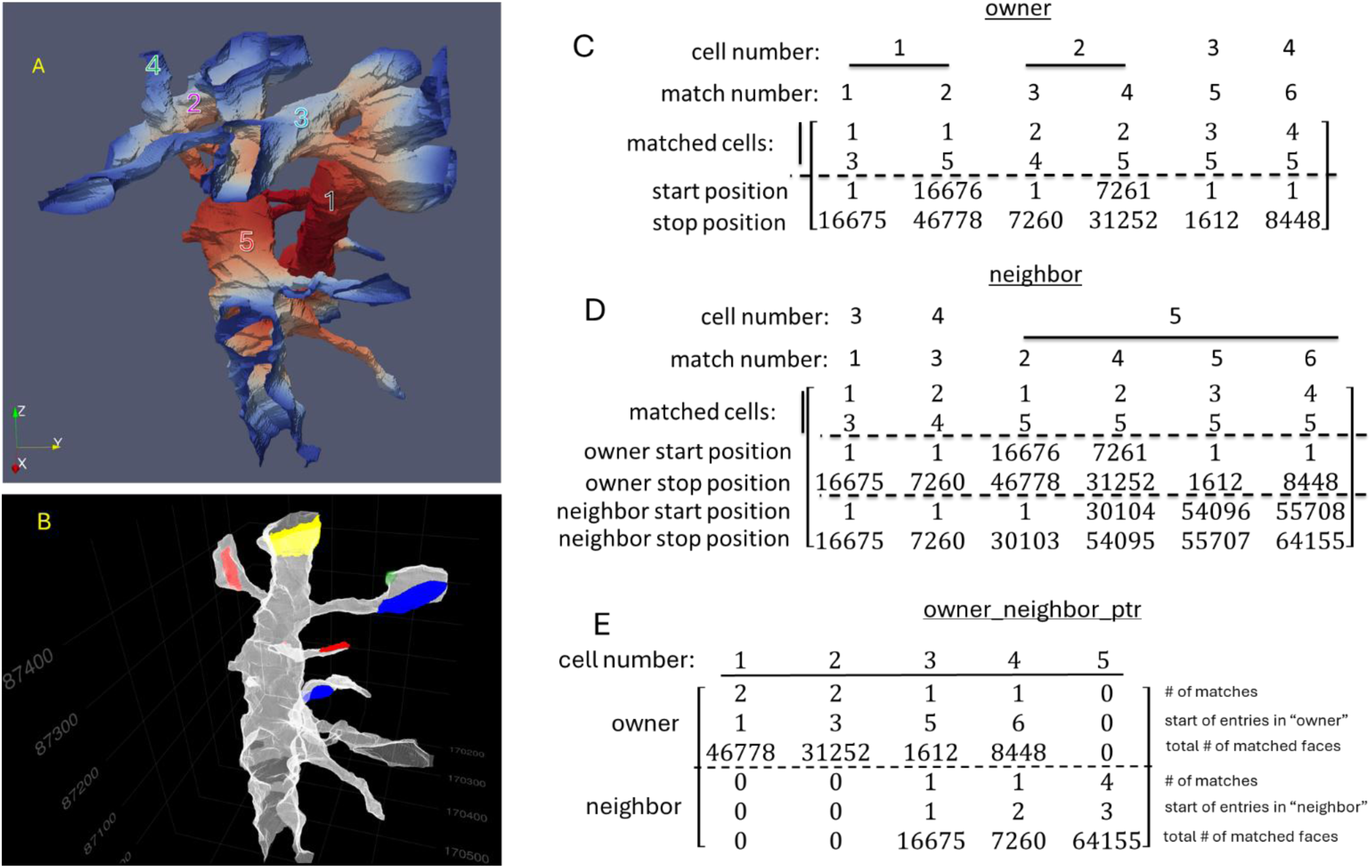
A) Five example cells that share faces within a 4 μm x 4 μm x 4 μm analysis volume. B) Shared faces between cell 5 and cells 1 (green), 2 (yellow), 3 (blue) and 4 (red). C) Matrix ‘owner’ records matches between cells. The first row are cells that own a match, and the second row are the corresponding neighbor cells. The third row is a pointer to the start of the list of matched faces in the vector ‘owner_match’ and the fourth row is a pointer to the end of the list. D) ‘neighbor’, similar to ‘owner’, but sorted by the second row. The third and fourth rows are the same as in owner, but the fifth and six rows are pointers to the vector of matched faces of the neighbor in a match, which are stored in the array ‘neighbor_match. E) The first three rows of ‘owner_neighbor_ptr’ provide a map between a given cell number and the location of the cell’s owned matches in ‘owner’. The last three rows provide the same map for ‘neighbor’.

**Figure 6:**
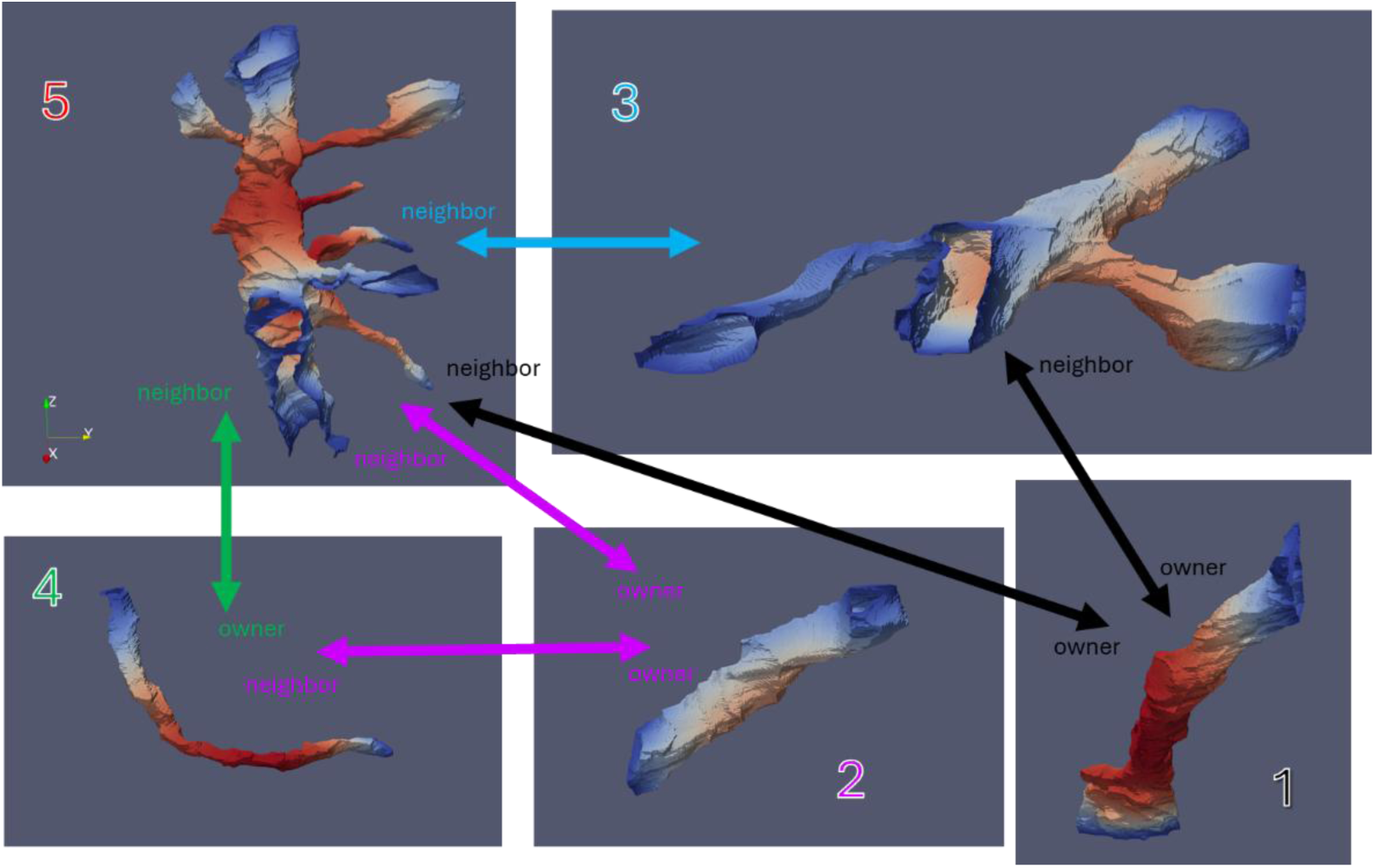
Matches between cells 1-5. The owner of a match is the cell with the lower cell number. Cell 1 owns matches with cells 3 and 5 (black). Cell 2 owns matches with cells 4 and 5. Cell 3 owns the match with cell 5. Cell 4 owns the match with cell 5. Cell 5 owns no matches but is a neighbor to cells 1-4.

**Figure 7:**
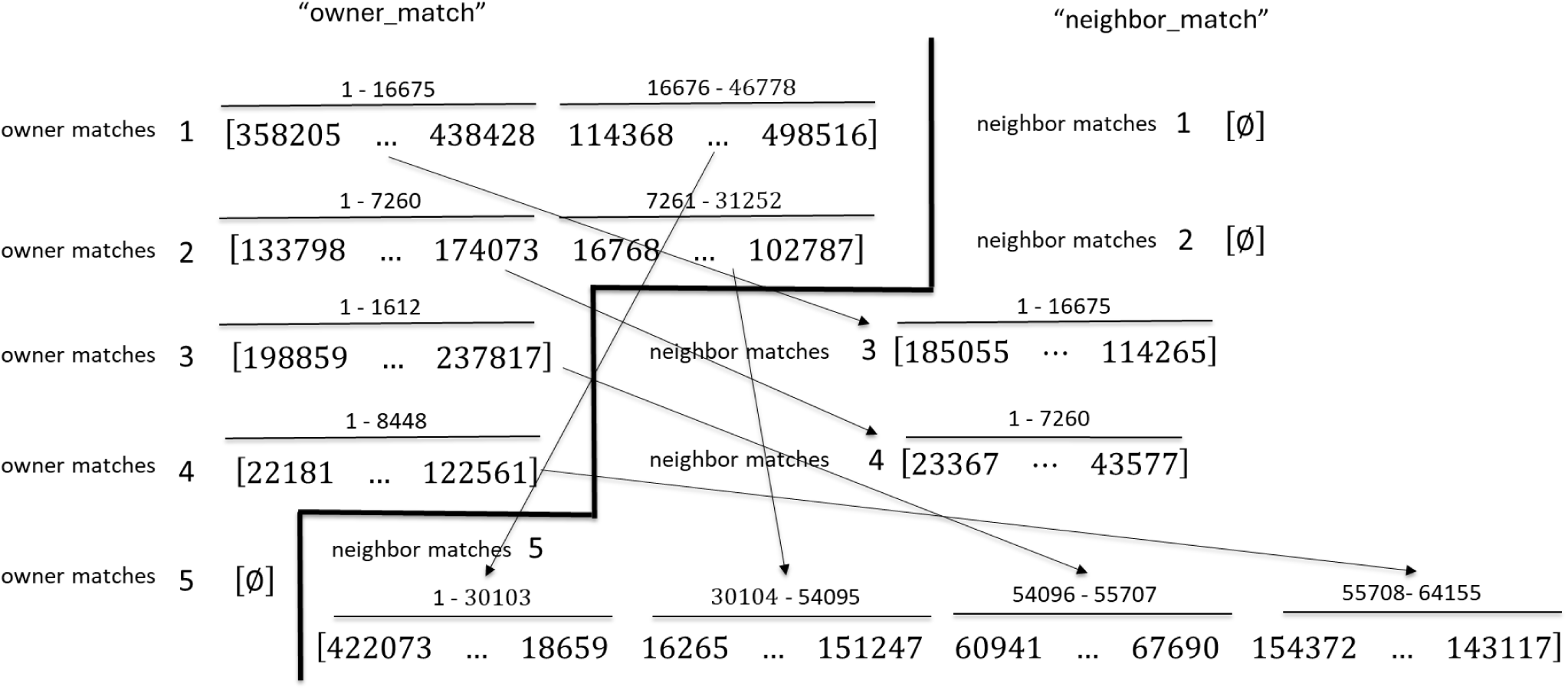
An illustration of the list of face numbers involved in matches. The order of the list of matched faces is the same in ‘owner_match’ and ‘neighbor_match’ wherever a match is present. Note that each cell stores all its matched faces in a single vector, and the start and end positions of the face number list for each match is found using the arrays ‘owner_neighbor_ptr’ and ‘owner’ or ‘neighbor”. Arrows illustrate matched lists of faces.

The owner-neighbor relationships for the five cells are shown in Figure 6. The face storage vectors ‘owner_match’ and ‘neighbor_match’ are shown in Figure 7. Each cell uses one vector for its list of owned matches and one vector for its list of neighbor matches (if there are no matches it is a single element null vector). Arrows show the correspondence between entries in ‘owner_match’ and ‘neighbor_match’.

Additionally, above each vector is the size of the entry, which can be compared to the values in ‘owner’ and ‘neighbor’. The matched faces are in the same order in ‘owner_match’ and ‘neighbor_match’ since that the vectors were sorted by the position of the centroid when created. Figure 8 shows that after a solution has been calculated for a cell, the concentrations at all faces in ‘owner_match’ are stored in ‘phi_owner’, and concentrations at all faces in ‘neighbor_match’ are stored in ‘phi_neighbor’. The corresponding face concentrations are averaged by the manager and then stored back into ‘phi_owner’ and ‘phi_neighbor’.

**Figure 8:**
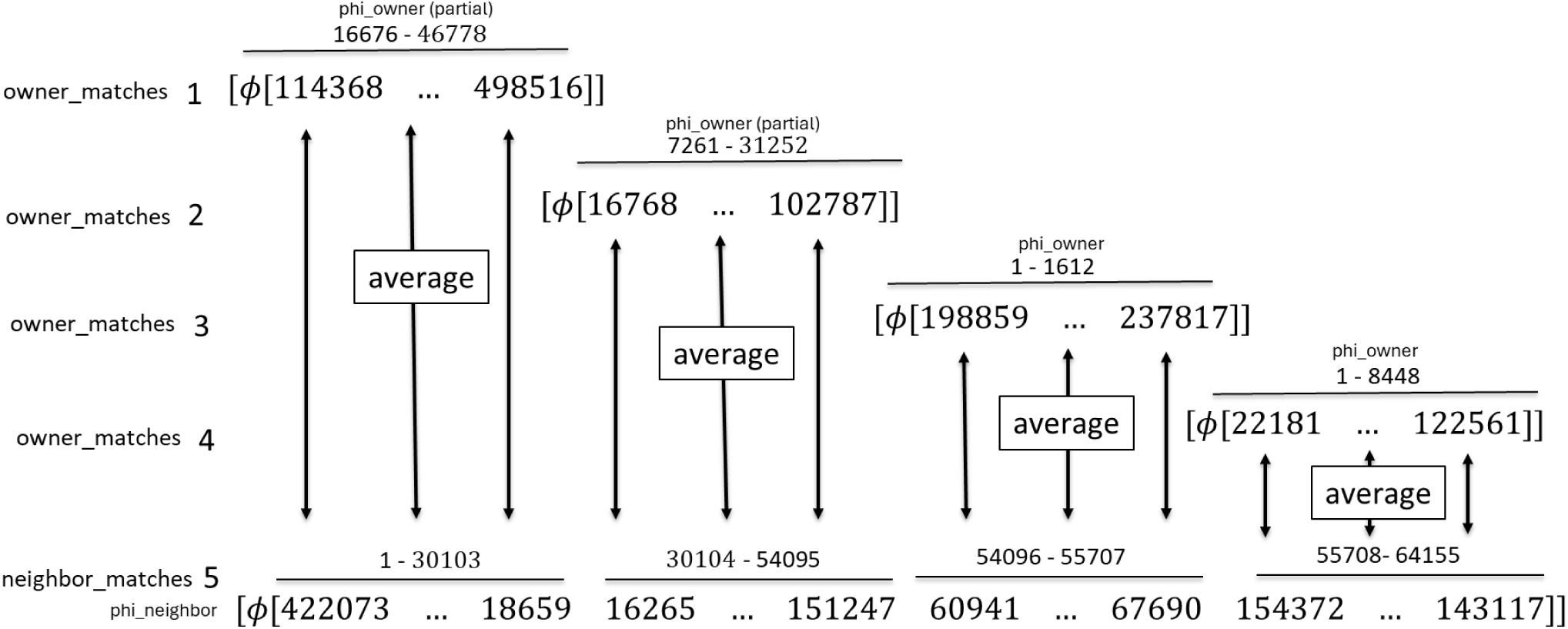
The two lists of matched faces are used to query the concentration (φ) vectors that store the current concentrations for all faces on the cell. The retrieved concentrations are stored in float32 vectors “phi_owner” and “phi_neighbor”. The notation φ[…] refers to the list of concentrations for the given face list.

The overall process is executed by a job scheduler as shown in Figure 9. At the start of each time point, the manager core will assign a biological cell to a worker. The worker will solve the linearized equations to find the concentrations at that time step (Figure 9: Step 1). When finished, the worker writes the solution at all faces to a vector ‘phi’ specific for that cell, stored on the manager node using one-sided RMA communication. The worker also writes concentrations into ‘phi_owner’ and ‘phi_neighbor’, also on the manager node. After all cells in the analysis volume have been processed, the manager then averages the concentrations between matched faces and writes the results back in the ‘phi_owner’ and ‘phi_neighbor’ vectors (Step 2). The manager then spins up more workers that read in the updated ‘phi_owner’ and ‘phi_neighbor’ vectors (Step 3). The workers update the ‘phi’ vector with the new values and write it to disk.

**Figure 9:**
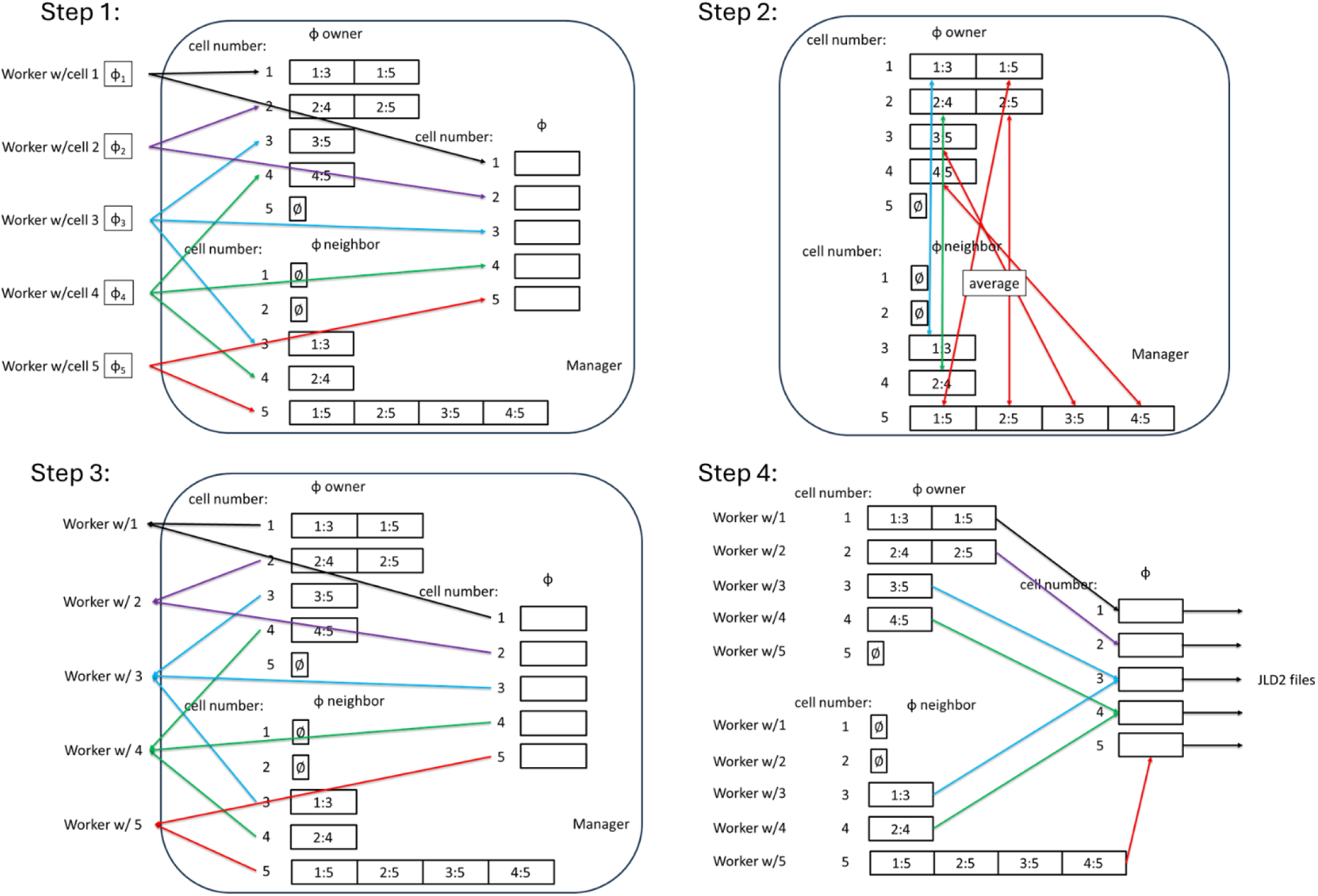
Protocol for mass transfer across matching faces using a manager/worker job scheduler. Step 1: After using the conjugate gradient method to solve for concentrations (ϕ) at each time step, the concentrations on all faces of a cell are recorded in the array of vectors ϕ on the manager node by the workers using one-sided communication. Concentrations on shared faces are also stored by the workers in ϕ_owner_ and ϕ_neighbor_ on the manager node. Step 2: The average value of each pair of faces in ϕ_owner_ and ϕ_neighbor_ is calculated and written back into both vectors by the manager. Step 3: The manager launches new workers that read the values of ϕ, ϕ_owner_ and ϕ_neighbor_ via one-sided communication. Step 4: Each worker writes updated ϕ_owner_ and ϕ_neighbor_ values into ϕ, which is then saved in a JLD2 file that is used in the next time step.

### Arrays of analysis volumes

Matches only occur within analysis volumes. However, the biological cells extend across boundaries. Edges of faces may lie on the faces of analysis volumes but never cross the boundaries. Each boundary edge is associated with one face. If there is another analysis volume adjacent, boundary edges will be shared by faces on opposite sides of the boundary. These faces must transfer mass across the edge if a concentration gradient is present. Thus, the concentrations of the faces immediately across the boundary must be known. Fortunately, the faces that share a boundary edge are part of the same biological cell, so the problem is somewhat limited in scope if there is a map of how each 18-digit identifier is translated into cell numbers in the different volumes.

Figure 10A shows the same cells as in Figure 5 but within an array of eight analysis volumes. The octant corresponding to Figure 5 is shown as a red-to-blue heatmap of concentrations, while the portions of the cells in other octants are shown in solid colors. Figure 10B shows the array of analysis volumes, with the volume in Figure 5 circled and highlighted.

**Figure 10:**
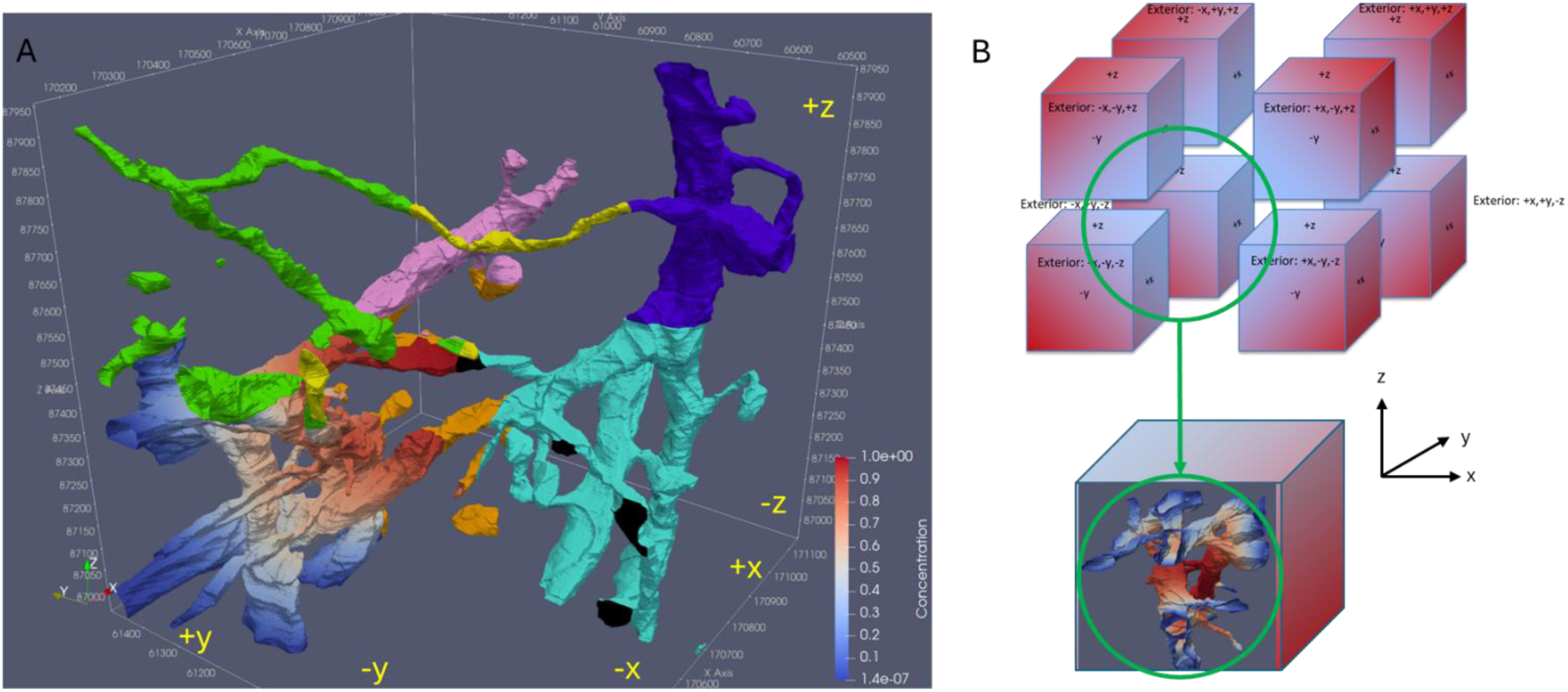
A. Illustration of the five cells from Figure 5 extending into neighboring analysis volumes. The original analysis volume is a heatmap of the concentrations at an intermediate time point. Neighbor volumes in the +x, -y and +z directions are shown in solid colors. B. The original analysis volume is in the lower back left in this and subsequent figures.

The face numbers of the boundary faces are stored within the array ‘interior_boundary_faces’, which is stored for each cell and is derived from the ‘boundary_edges’ array (Figure 11A). The analysis volume from Figure 5 is ‘Cube 3’, while the analysis volume in the positive z-direction is ‘Cube 4’ (analysis volumes will be called ‘cubes’ here). The faces of the analysis cubes are numbered 1-6, with -x face= 1, +x face = 2, -y face = 3, +y face = 4, -z face = 5, +z face = 6. Cube 3’s ‘cube face 6’ is Cube 4’s ‘cube face 5’. Any edge in Cube 3 that lies on ‘cube face 6’ will overlap with an edge on ‘cube face 5’ of Cube 4. Furthermore, due to the marching directions in the Marching Cubes algorithms, the order of the edges and thus faces in ‘interior_boundary_faces’ is the same on both sides of a boundary without further intervention. It is thus not necessary to exchange face numbers between analysis cubes. The relationship between a boundary edge, adjoining faces and analysis volumes is illustrated in Figure 11B-D. The cell number in cube 4 will be different from the cell number in cube 3, and the cell numbers will appear in the lists in different orders.

**Figure 11:**
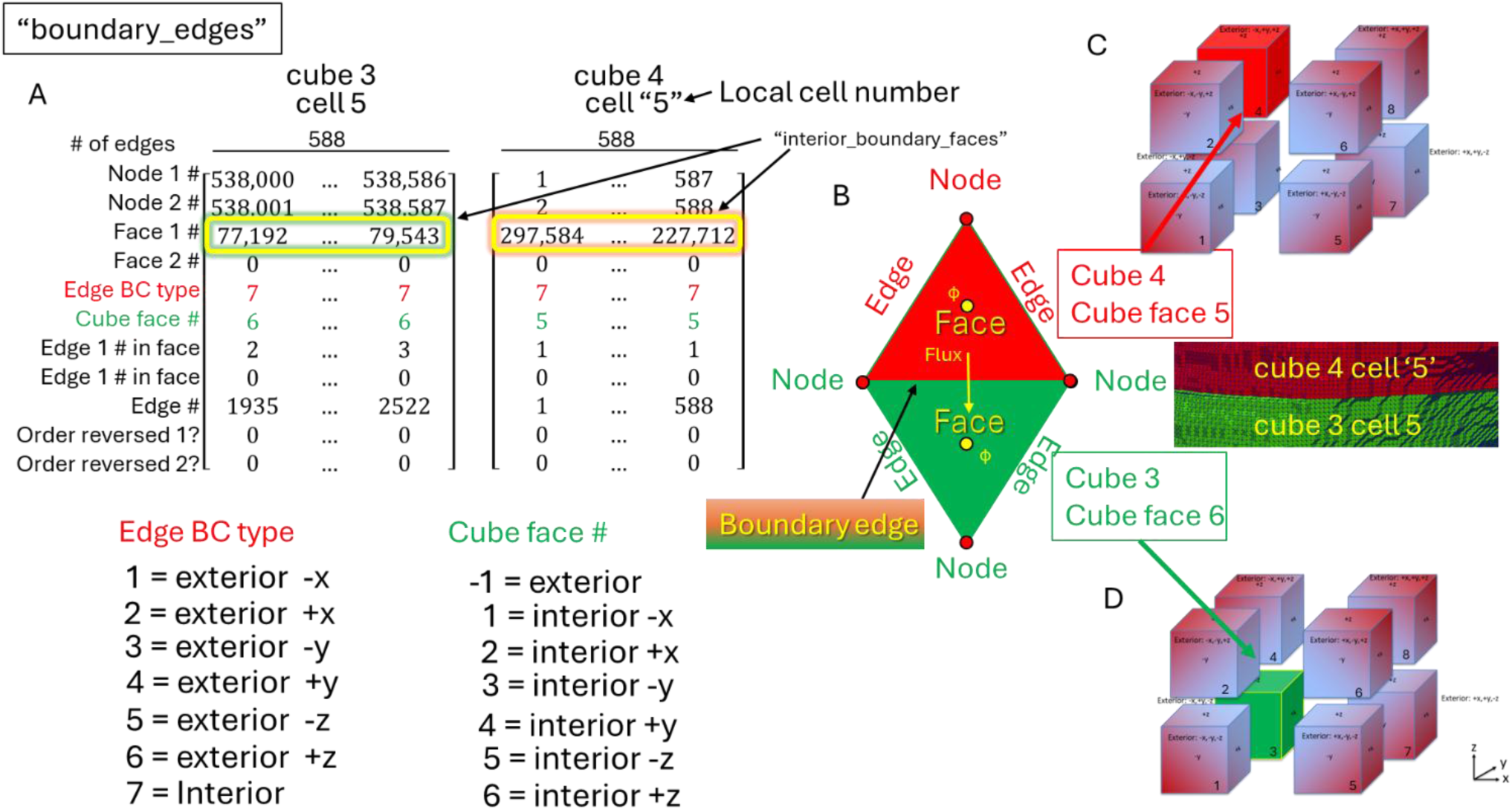
Edges on the faces of analysis volumes. A. Edges that are on analysis volume faces are recorded in the array ‘boundary_edges’, which is a modified form of ‘edges’. The Edge BC type is set to 7 if the boundary edge is on an interior boundary and set to 1-6 if on an exterior boundary. If the boundary edge is on an interior boundary, the edge number of the boundary edge in the neighboring analysis volume is written into ‘Edge #’. B. An example interior boundary edge. The red analysis cube (cube 4) is in the +z direction from the green analysis volume (cube 3). The boundary faces must exchange concentration values at each time step so that the flux across the boundary may be estimated in the following time step. C. Cube 4 (red) in relation to other analysis volumes. D) Cube 3 (green) in relation to other analysis volumes.

Thus, it is necessary in the setup program to map the cell numbers and point to the proper entry point of face numbers in ‘interior_boundary_faces’. The number of entries will be the same; in this example there are 588 boundary edges for this cell on each boundary.

The pointers to the list of face numbers are stored in an array ‘cells_on_faces’, where columns are cell numbers (Figure 12). The first six rows store the total number of boundary edges. A negative number signifies a cube face that is exterior to the array of analysis volumes. No mass is transferred at these cube faces, but boundary conditions must be applied. Cube faces that are interior to the array of analysis volumes have a positive number of edges. Notice that column 5/row 6 of ‘cells_on_faces’ for cube 3 corresponds to the example from Figure 11. Rows 7-12 store pointers to the start of the list of boundary faces in the vector ‘interior_boundary_faces’. The vector ‘interior_boundary_faces’ is cell-centric, because each cell carries its own list of interior boundary faces. This is efficient because each cell only needs to know its own face numbers on the boundary, which the worker uses to query the concentration values for the boundary faces immediately after solving the linearized equations. Rows 12-18 contain pointers into a vector that stores the concentrations of the interior boundary faces. The concentration list is stored in a cube face-centric manner, meaning that each face curates the list of concentrations for all the cells on the face. This allows for an efficient exchange of concentrations at the end of each time step because the entire cube face may be analyzed in one step.

**Figure 12:**
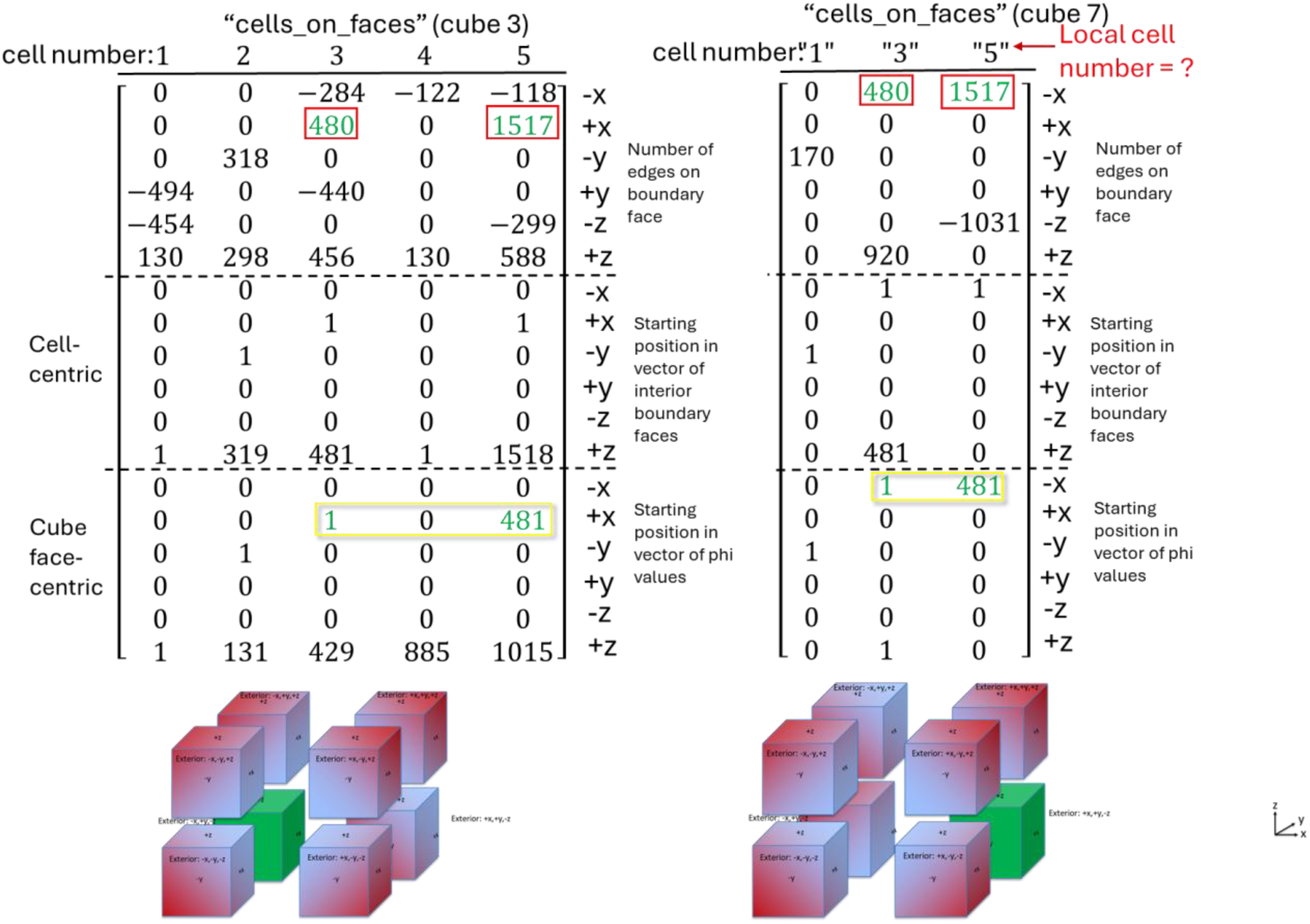
The table “cells_on_faces” is used to point to the starting locations in the list of faces. Each column corresponds to a unique cell. The first six rows store the number of boundary edges for a cell on each of the six faces of the analysis volume cube. A negative number of edges means that the face is an exterior face. The middle six rows are ‘cell-centric’ meaning that all boundary faces for the cell are stored in one array, while the bottom six rows are ‘cube face-centric’ because the boundary concentrations for the whole face are stored in one vector. The cell numbers for cube 7 are listed by their cell number in cube 3, but the corresponding cell numbers in cube 7 will be different and in a different order. The ‘cell-centric’ values are used to find faces in the vector ‘interior_boundary_faces’. The ‘cube face-centric’ values are used to find face concentrations.

The cube face list of boundary face concentrations is stored by the manager using one-sided RMA communication (Figure 13). Each analysis cube is solved separately and sequentially, and the lists of cube face concentrations is stored in ϕ_conc_ (‘phi_conc’), where ϕ symbolizes concentration. The 588 boundary face concentrations from Figure 11 are the last entry of ϕ_conc_ for cube face 6 (Figure 13A – red circle).

**Figure 13:**
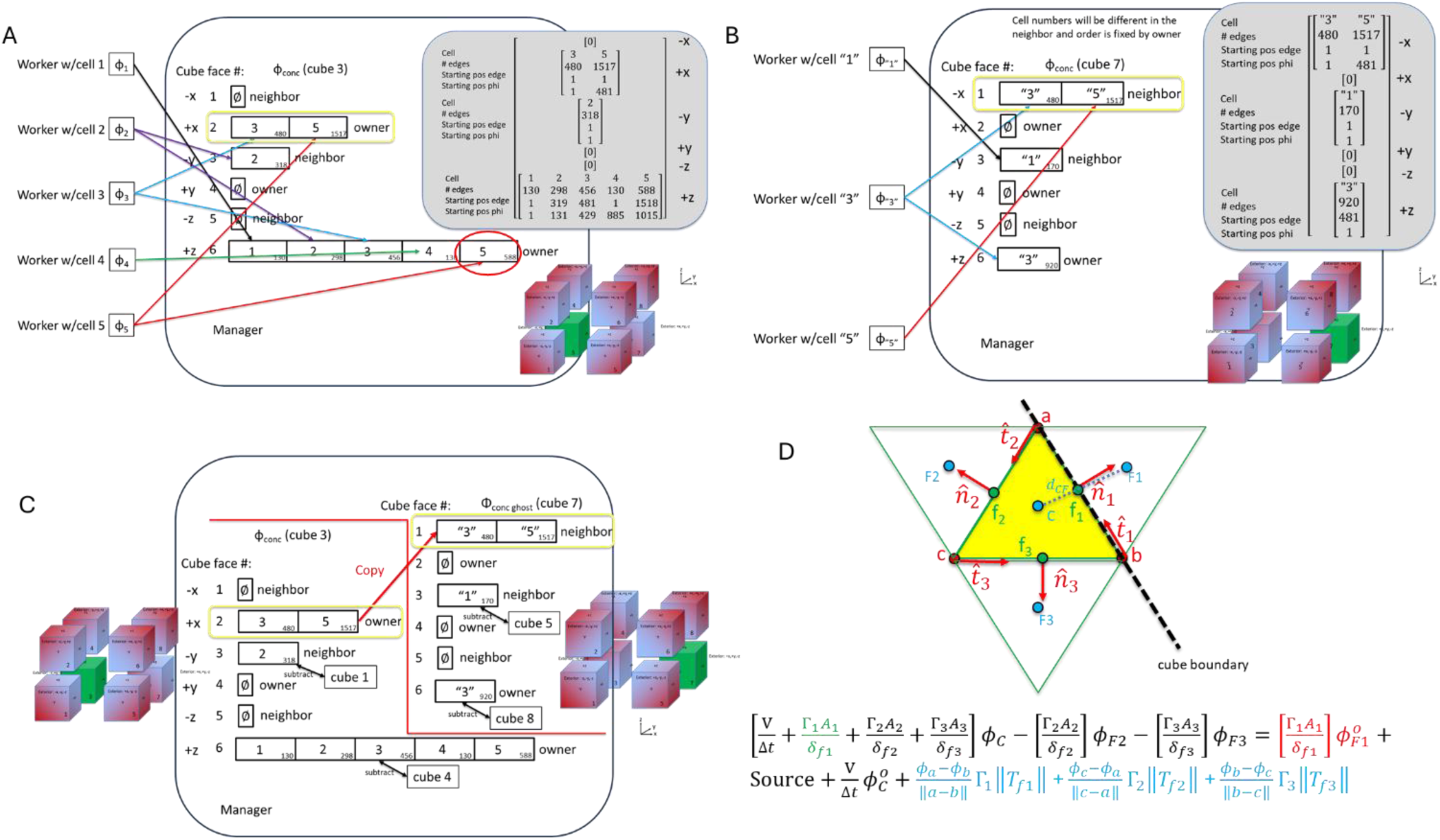
Protocol for mass transfer across adjacent analysis volumes. Biological cells that span analysis volumes exchange mass across the boundary at each time step based on the concentration gradient between adjoining faces. A. For cube 3, a worker solves for ϕ on all faces of a cell, then reads the list of interior boundary faces and writes the concentrations from those faces into ϕ_conc_. B. A similar procedure is followed for cube 7. C. At the end of each time step, the manager writes ϕ_conc_ into ϕ_ghost_ of its neighbor cube. D. At the next time step, ϕ_ghost_ will be used to estimate the mass flux at the boundary using a semi-implicit approach, with the implicit portion in green and the explicit portion in red. The cyan term is the non-orthogonality correction.

Because cell 5 also passes through cube face 2, the worker handling cell 5 also writes the appropriate concentrations starting at position 481 of ϕ_conc_ for cube face 2 (Figure 13A - yellow rectangle). The worker with cell 3 also writes into the same ϕ_conc_ vector for cube face 2, but starting at position 1. When cube 7 is analyzed (the analysis cube in the +x direction from cube 3), the workers with cells ‘3’ and ‘5’ write the appropriate concentrations in the vector ϕ_conc_ for cube face 1 (cube face 1 because the relevant boundary edges are on cube face 1 of analysis cube 7) (Figure 13B – yellow rectangle). Then, the manager will write ϕ_conc_ from cube 3 into a vector ϕ_ghost_ for cube 7 and write ϕ_conc_ from cube 7 into a vector ϕ_ghost_ for cube 3 (Figure 13C). One could simply overwrite the concentrations; however, this will lead to failures if the simulation is halted during the transfer. To deal with restarts, a time step is only marked as complete after the ghost concentrations are written. Otherwise, the simulation will restart from the previous time step.

The list of concentrations is stored in two locations, one for odd time steps and the other for even time steps, allowing access to the last completed time step in all cases.

The cube boundary is depicted as a dashed black line in Figure 13D. The yellow face needs the concentration ϕ_F1_ to calculate the explicit part of the flux at the next time step according to the equation shown. The equation includes a non-orthogonal correction. Terms on the left-hand side may be calculated ahead of time. Terms on the right hand side are updated at each time step (explicit part of the flux, source term, non-homogenous part of the time difference) or at each iteration (non-orthogonal correction). The non-orthogonal correction here uses the nodal concentrations at a, b and c to approximate the gradient tangential to the face by a first order difference equation.

## Discussion

The primary result of the work is a framework that allows solving the convection-diffusion equations on thousands of objects in a parallel manner. Results generated from the framework with biological cells are demonstrated in a companion publication. The thousands of objects that are analyzed are in contact and exchange mass, which requires bookkeeping to record the matches. This is handled using a manager-worker model with one-sided RMA via MPI to minimize duplication of arrays and to allow the manager to perform summary calculations. Most of the thousands of objects also pass through the boundaries of analysis volumes. Each analysis volume is solved sequentially during a time step, with the important results again stored on the manager core and communicated to and from the workers by one sided RMA via MPI. The manager persists throughout the entire analysis period, switching from equation solution to face matching to concentration bookkeeping within each time step.

The current approach is unique in the attempt to model mass transfer in thousands of interacting objects.

The strengths are that the simulation of 2175 neurons and other brain cells at 8 nm resolution in an 8 μm x 8 μm x 8 μm volume is feasible with only a single node with 40 cores in a reasonable time, as shown in the companion publication. The weaknesses of the approach stem largely from limitations of the incoming data: 1) the extracellular space is poorly defined in the source images, 2) segmentations of the source images contain errors leading to high curvature structures that are non-physical and harmful to accuracy, and 3) resolution in the z-direction is 5x worse than in the x and y directions. To introduce volume to the extracellular space, the surface meshes may be ‘inflated’ to produce the 3D space. However, the quality of the inflated elements will depend a great deal on the smoothness of the surface mesh. While the surface mesh smoothing and interpolation algorithms used here lead to much smoother meshes, additional smoothing will be extremely beneficial. The current mesh smoothing and interpolation procedures are described in a second companion publication. Additional smoothing will likely come from Taubin smoothing algorithms,^15^ however, these methods must be expanded to handle hundreds to thousands of objects simultaneously while preserving the interfaces between the objects. This is the topic of ongoing work.

Solution of the convection-diffusion equation on surface meshes is an approximate method. For convection, the velocities along the mesh must be known, which would involve solution of the Navier-Stokes equation. If the velocity is perfectly aligned with the surface mesh, then solution of the convection-diffusion is possible on a 3D surface mesh. This can be done with some model geometries as shown here. This is not applicable to the biological cell geometries.

Solution of the diffusion equation on surface meshes is sometimes used to determine geodesic distances on the surfaces of complex objects by the ‘heat method’.^8,9^ The accuracy of the solutions will depend on the smoothness of the mesh. In the heat method, the magnitudes of temperature gradients are irrelevant, only the gradient directions are required for the method. Here, the accuracy suffers due to approximations in estimating the areas of the faces of the corresponding volumetric mesh and differences between centroids of triangular faces and the centroids of the implied prismatic volumes of the corresponding volumetric mesh. When curvature is zero, the equations are highly accurate as shown. As curvature increases, accuracy suffers from loss of mass balance due to the approximations. The reason these limitations were accepted was because advancements in mesh smoothing algorithms will be more impactful on accuracy than improvements to the discretization.

The Julia language has become an important language for scientific computing. The Julia language was designed to achieve C-like speeds while using Python- or MATLAB-like syntax. In practice, C-like speed is achieved by minimizing allocations and avoiding type instability. Python and MATLAB have improved their speeds largely by writing computationally intensive code in compiled languages like C or Fortran. In Julia, C-like speed can be achieved with native Julia code by following a few simple performance guidelines that produce code that is fast on its own but also easily converted to a compiled language. This makes Julia an excellent candidate for a rapid-prototyping language.^16^ A group at Meta developed a new audio protocols that was developed in Julia and deployed in C/C++ (https://youtu.be/3ypsZUNRjI4?t=494). In that same spirit, it is advantageous in the current project, where multiple approaches need to be developed, to take advantage of Julia’s ease of use and package management to rapidly develop new solutions to the current ‘thousand object’ CFD problem.

Handling thousands of cells as individual objects is an interesting question, but it is important to remember that once volumetric meshes are developed for this problem, it may not make sense to treat individual cells as unique objects. Even before that step, fusing the surface meshes into one object per volume is also a valid approach. The only potential weaknesses are that the stability of the solution methods might suffer, and the size of the matrices will increase dramatically. In the current approach, there are 2-5 cells out of 2145 that are small and near boundaries and whose solutions are unstable. As such, solutions for cells are tested for any undefined values and if present, are set to zero. It is difficult to predict if such structures would lead to difficulties with fused meshes. One can also imagine that it may be easier to incorporate electrophysiology, effects of myelination and diurnal effects in the model if cells are treated as unique objects. Answering such questions may open the door not only to addressing questions about the clearance of toxins from the brain but may facilitate large scale models of brain computations.

The future challenges to be addressed include: 1) Further smoothing of surface meshes, 2) Inflating surface meshes to produce volumetric meshes (with mesh fusion or with individual cells), 3) Adding pressure-velocity coupling to the volumetric meshes, 4) Expanding from 8 analysis volumes to 32,768 to capture a capillary bed, 5) Defining boundary conditions at capillaries to allow pressure and flow fluctuations. Even after adding those capabilities, it will still remain to include the effects of volume changes within the blood vessels due to pressure waves, the potential for changes in axon and dendrite diameters due to external and internal factors, the effects of astrocytes on fluid transport due to their aquaporin channels, and dynamic effects such as remodeling of synapses and removal of plaques and tangles by microglia. The current work is a first step down this path.

## Conclusions

Existing computational tools are not well-suited to automating the analysis of thousands of cells in a tissue, particularly convection and diffusion of solutes in the narrow gap between these cells. The current work shows a framework for an analysis of diffusion around cells in the brain cortex. The framework parallelizes by biological cell and handles mass transfer between the cells at each time step, as well as mass transfer at analysis volume boundaries. This method is a first step towards developing high resolution simulations of mass transport in the brain at TEM-level resolution.

## Acknowledgements

The work was supported by startup funds at the University of Washington

## Author contributions

DLE designed the study, designed the framework, wrote code and wrote the paper

DPE aided with design of the study, provided feedback on the framework, wrote code and edited the paper.

## Data availability

Programs used to generate the results are available at: https://github.com/elbert5770/neuron_transport_3D_v3_archive.

